# Structural and Functional Characterization of Novel Phosphotyrosine Phosphatase Protein from *Drosophila Melanogaster* (Pupal Retina)

**DOI:** 10.1101/2022.02.19.481152

**Authors:** Rubina Naz, Anwar Iqbal, Asma Saeed, Alamzeb Khan, Meshal Shutaywi, Adriana Lavinia Cioca, Narcisa Vrinceanu

## Abstract

A novel pair of protein Tyrosine Phosphatases in *Drosophila Melanogaster* (pupal retina) has been identified. Phosphotyrosyl protein phosphatases (PTPs) are structurally diverse enzymes increasingly recognized having fundamental role in cellular processes including effects on metabolism, cell proliferation and differentiation. This study presents comparative homology modeling of low molecular weight phosphotyrosine protein phosphatase (PTPs) from *Drosophila melanogaster* (Dr-PTPs) and their complexation with potent inhibitor HEPES. The 3D structure was predicted based on sequence homology with bovine heart low molecular weight PTPs (Bh-PTPs). The sequence homology is approximately 50% identical to each other and to low molecular weight protein tyrosine phosphatases (PTPs) in other species. Comparison of the 3D structures of Bh-PTPs and Dr-PTPs (primo-2) reveals a remarkable similarity having a four stranded central parallel β sheet with flanking α helices on both sides, showing two right-handed β-α-β motifs. The inhibitor shows similar binding features as seen in other PTPs. The study also highlights the key catalytic residues important for target recognition and PTPs activation. The structure guided studies of both proteins clearly reveal a common mechanism of action, inhibitor binding at the active site and will expected to contribute towards the basic understanding of functional association of this enzyme with other molecules.

## 1. Introduction

Low molecular weight phosphotyrosine protein phosphatases (PTPs), previously known as low molecular weight acid phosphatases, catalyzes the hydrolysis of tyrosine phosporylated proteins, low molecular weight aryl phosphates and natural and synthetic acyl phosphates ^1,2^. Although the activity of PTPs on serine and threonine phosphorylated proteins are very poor with the exception of flavin mononucleotide (FMN) ^3,4^. Tyrosine phosphorylation plays a vital role in the regulation of the variety of developmental processes. These processes include several cell functions like growth, cell differentiation, metabolism, cell cycle and cyto-skeletal functions. Furthermore, the phosphorylation state of tyrosine and seγ/thr of signaling proteins are controlled via specific reaction. Thus, the phosphorylation state is controlled by a very dynamic way to avoid severe malfunction of cell ^5^.

PTPs can act as tumor suppressor by inhibiting cell growth. Functionally two types of PTPs sequences are conserved and well distinguished by structure comparison. The one known as classical PTPs are specific for tyrosine residues and other with the dual-specificity phosphates (DSPS) are essential for serine and threonine dephosphorylation. The active site (C-(X)_5_-R) of these PTPs contains several conserved cysteine and arginine, important for catalyzing phosphorylation processes, and thus plays a vital role in regulation of signal transduction. All these acid phosphatase enzymes share little sequence homology, different range of molecular weight (18 kDa or above), but exhibit same catalytic mechanism ^6,7^. The structural features of low molecular weight PTPs comprises relatively different fold comparing from fission yeast to mammals ^8,9^. The overall three dimensional structural features contains four B sheets at center and surrounded by a helices ^6,10,11^. However, several similarities can be seen in structural features and binding side pockets of PTPs to Mr-PTPs ^10–15^. Importantly, the conserved the p-loop stabilized by complex hydrogen network and favor the phosphate trigonal bipyramidal transition state geometry ^16–19^. Thus all low and higher Mr-PTPs and PTPs shares identical catalytic mechanism ^20^. The enzymatic reaction is triggered by the first cysteine, where the substrate binding at active site is stabilized by the p-loop residues via hydrogen bonding and three anionic oxygen atoms. These transient interaction orient phosphorous atom in feasible position for nucleophilic attack and favors the enzymatic phosphorylation reaction ^20–22^. The nucleophilic attack of thiol (Sγ) group takes place in the presence of proton donor aspartic acid and thus phosphor-cysteine intermediate is formed ^23–25^. The formation of phosphor-cysteine intermediate is also favored and stabilized by the presence of p-loop where several residues are involved in binding and lowering the activation energy ^20^. In the subsequent step, hydrolysis of phosphor-enzyme intermediate complex via attack of water takesplace, resulting in the liberation of inorganic phosphate (Pi). The enzymatic phosphorylation via hydrolysis works well at wide range of pH 5.5-7.5 from substrate.

The structural details of proteins are important parameters to understand the reactivity and stability of proteins. Several advance techniques like X-ray crystallography, nuclear magnetic resonance (NMR) and Electron Microscopy are frequently used to determine the structure of proteins. However, theoretical approaches like comparative modeling often used as a useful alternative to other biophysical and analytical techniques by providing insights into structural and functional aspects of proteins. In the current project, the comparative modeling technique has been used for the 3D structural prediction of sequence emerging from Primo-2 of *Drosophila melanogaster* (fruit fly) classified as low molecular weight PTPs family (LMW-PTPs). The present work is designed to elaborate the prediction of evolutionary context of the sequence homological ancestors, structural aspects and active site conformational states of *Drosophila Melanogaster* PTPs (Dr-PTPs) and spatial geometry formation of the active site. Dr-PTPs shares 46% amino acid sequence identity with that of Bh-PTPs (PDB: 1DG9) particularly in active site regions.

## 2. Results and discussion

### 2.1. Sequence Analysis

Two novel pairs of proteins tyrosine phosphatases were identified in the drosophila pupil retina. It was found that the primary sequences were approximately 50%identical and characterized as low molecular weight protein phosphatase (Fig. 1&2). The first sequence was incorporated at primo-1 (155 amino acid) and the other encoded by primo-2 (164 amino acid).

**Figure 1.**
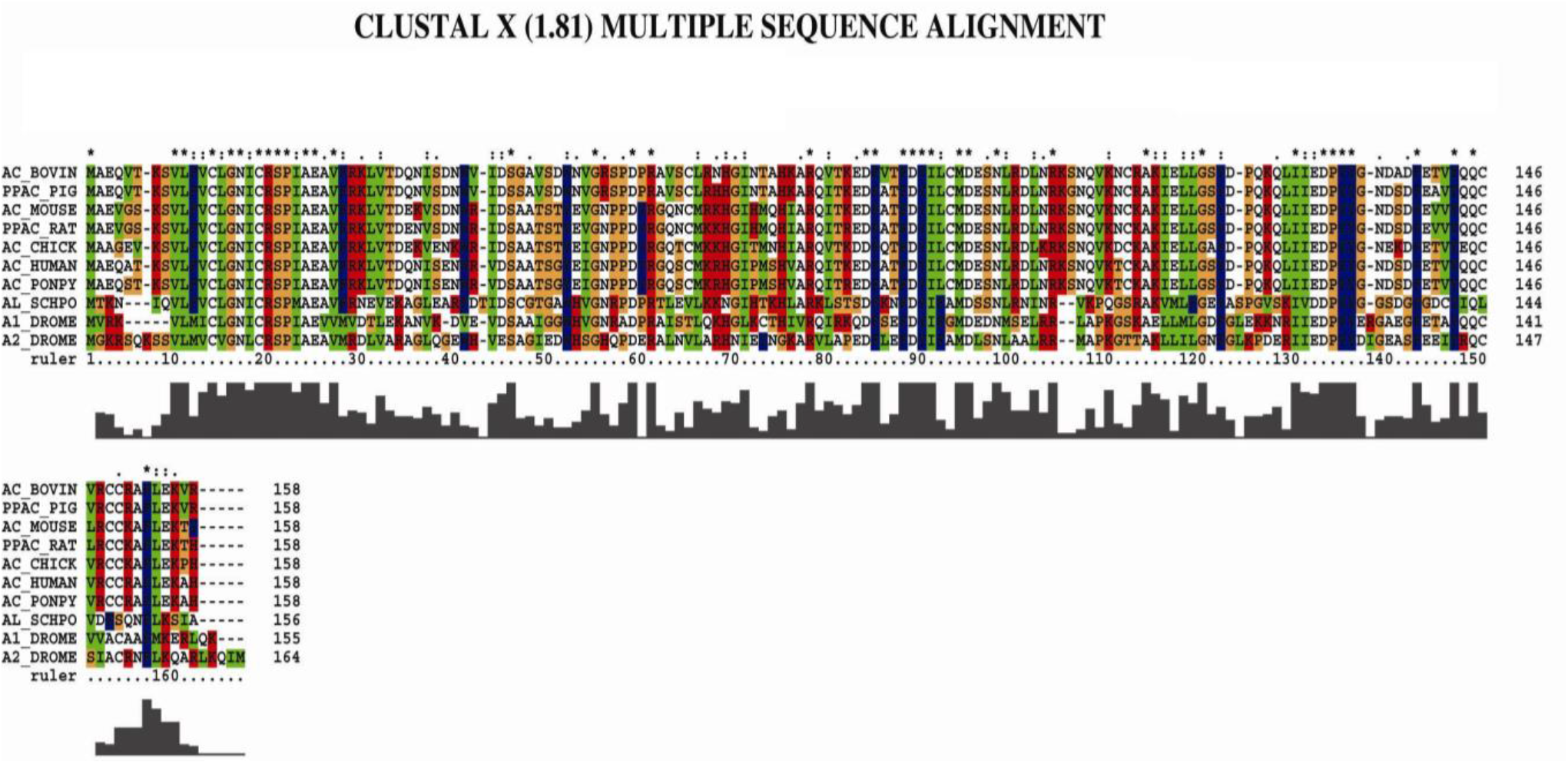
Multiple sequence alignment of 10 PTPs sequences. Sequences are named as Swiss-Prot entery first letter represents gene, the second part represents the biological source of gene. The symbol “*” represents strongly conserved “.” represents weakly conserved“:”represents identical residues

**Figure 2.**
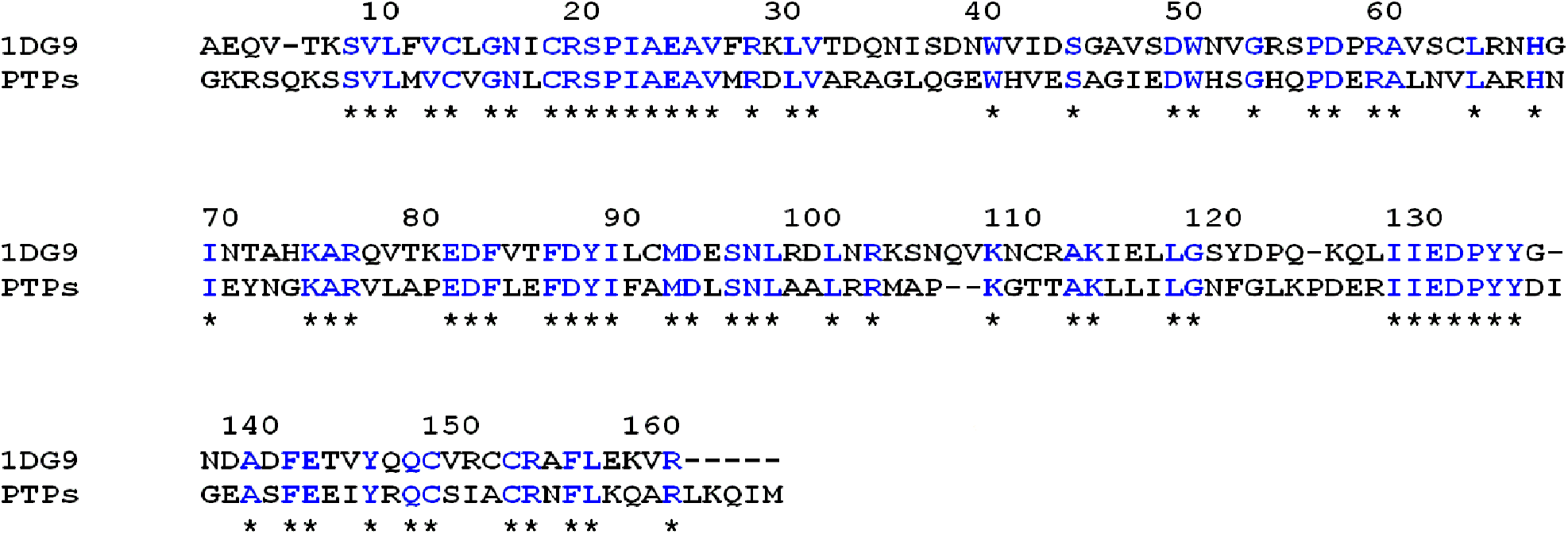
Pairwise sequence alignment used for building model of Drosophila phosphatase. Target sequence represented by PTPs. based on the structure of Bovin heart phosphatase template structure represented by1DG9.

### 2.2. Structure Topology

The structure of Dr-PTPs (primo-2) comprises a fold containing four central β parallel sheets gathered by a-helices: a right-handed β-α-β motif. The conserved sequence known as active site C—(X)_5_—R(S/T) of acid phosphatases was present as sequence CVGNLCRS in Dr-PTPs. The active site was present in the form of loop extending from between β 1 and helix α1 (Fig. 4).

The crystal structure (PDB: IDG9) shows a complex of Bh-PTPs with HEPES, where the active site residues were interacting with the sulfonate moiety, accumulating phosphate binding manner in the pocket. All the oxygen atoms of sulfonate group was involved in complex hydrogen bonding network, offered by N, H and S of the backbone residues of the active site loop and conserved arginine (Arg 18). The structural features of sulfonate complex were found similar to phosphates bounded at active sites. In our case, the highest sequence similarity of Dr-PTPs (Primo-2) primary sequence at substrate attracting site C-(X5)-R, tyrosine phosphorylation site RIIEDPYY was found. The 3D structure of Dr-PTPs (Primo-2) was superimposed at Bh-PTPs (Pdb: 1DG9). It was found the both structures aligned well for several motifs. However, regions like Gly 1, Lys 6, Pro 105, Thr 108, Ile 134, Glu 136, Lys 121, Asp 123 and Leu 159, Met 163 of the Dr-PTPs structural orientation was different from the target. Furthermore, the sequence analysis on the basis of multiple sequence alignment and the construction of phylogenetic tree (PHYLIP package) shows that both Dr-PTPs and Bh-PTPs belong to common ancestors. They are homologous sequences and members of same subfamily as per evolutionary context (Fig. 4).

### 2.3. Structure Topology

The overall secondary and tertiary structure of the Drosophila phosphatase strongly resembles to templates 1BVH, 1DG9, 1PNT (Bovin heart) and other low molecular weight phosphotyrosine proteins phosphatase (Fig 3). The 3D structure was characterized with the active site end at the α1 and situated close to the N terminal region followed by the P-loop and β1 strand. The active site emerged as deep groove encompassing aromatic residues ((Trp-49, Tyr-131, Tyr-132) appears like claws while the conserved residues Asp-56 and Arg 58 of variable loop provides a network of hydrogen bonds with other adjacent α5-helice and β4-strand. All these residues work together for the target recognition and sets a proper orientation of ligand for catalytically important residues Cys12, Cys17 and Arg18, whether Asp129 (β4 extend loop) on the other end of the pocket to facilitate the protonation of phosphorylated intermediate together with Tyr131 and Tyr132. The hydrophobicity of active site groove is maintained by several buried hydrophobic residues like Leu 9, Val11, Phe82, Ile 88, Leu 99 and Lys 102.

**Figure 3.**
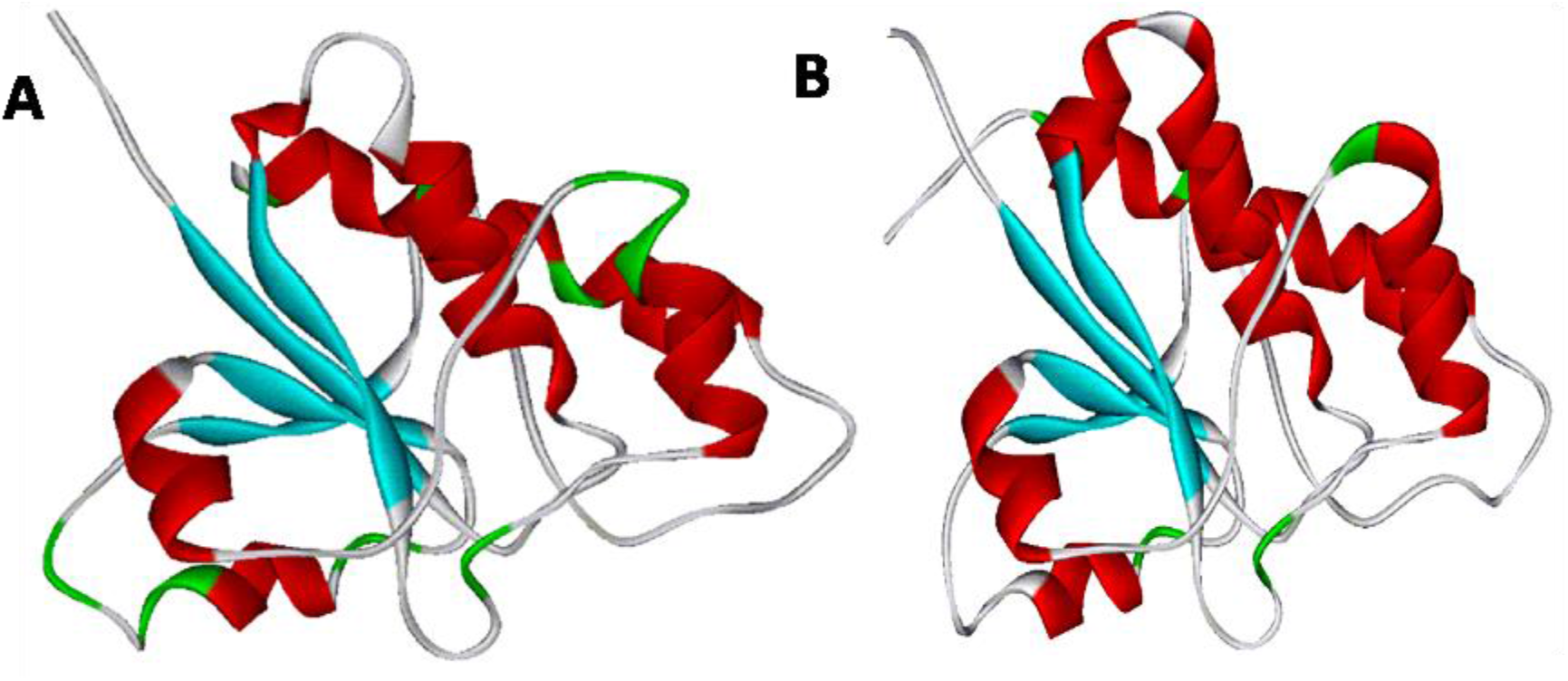
A complete Ribbon diagram of low molecular weight phosphotyrosine protein phosphatase. Showing α/β Proteins, α helix (Red), β Sheets (blue).(A) 1BVH(template), (B) Drosophila(target).

**Figure 4.**
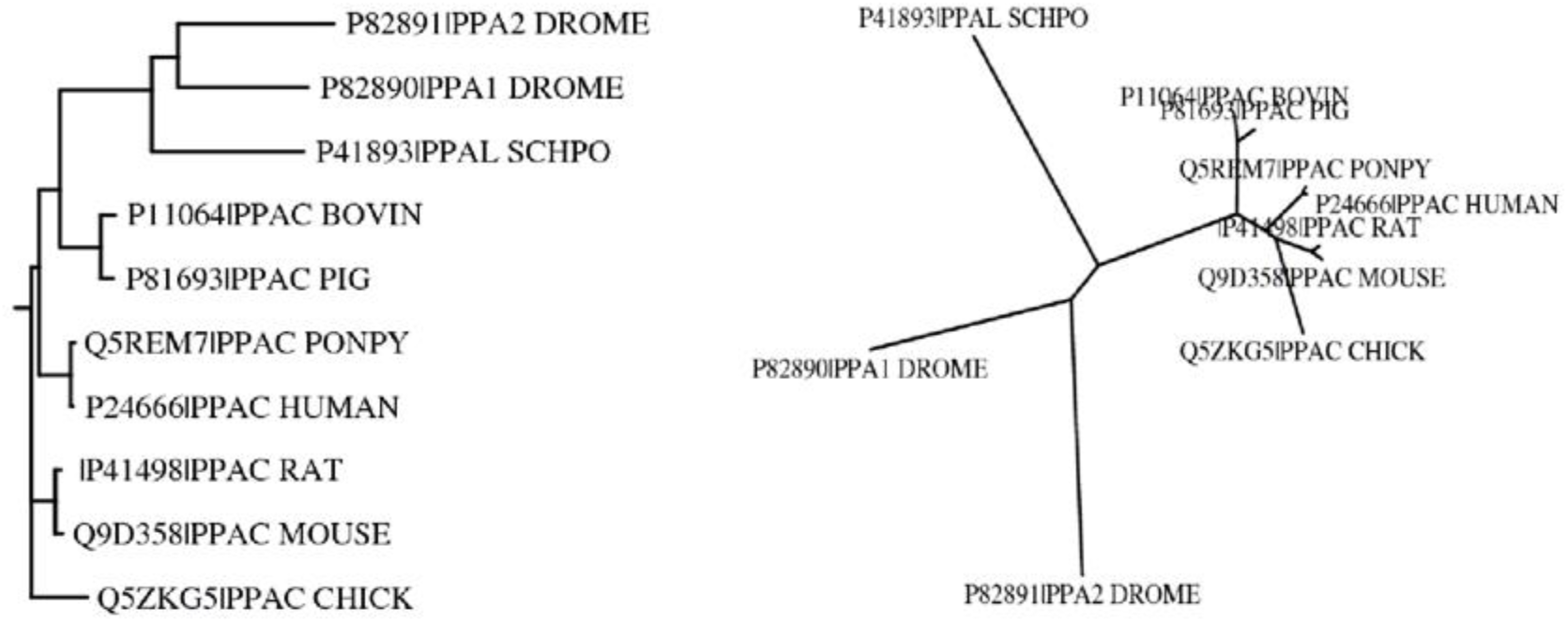
Dendrogram of PPA2 *Drosophila Melanogaster* and related proteins made by PHYLIP & depicts relationship among various forms of PPAC,PPAI, PPA2 genes. The first part of the code represents gene where as the second part of the code represents the source of protein.

**Figure 5.**
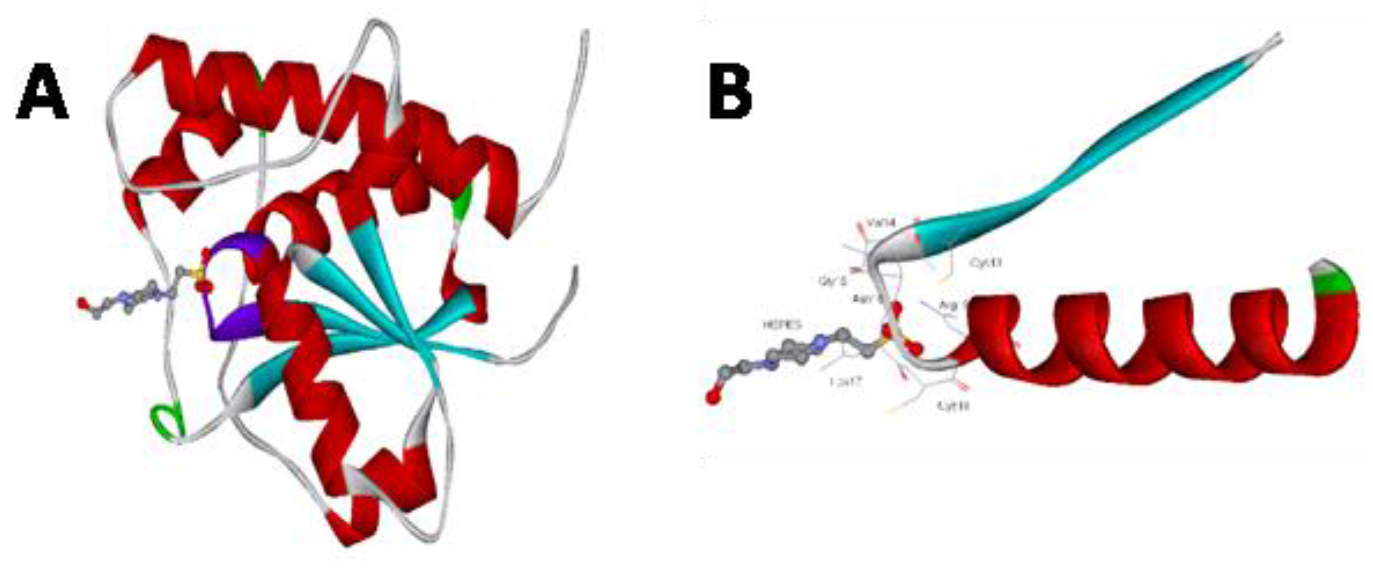
(A). Schematic diagram of Bovin heart PTPs (1DG9)complexed with HEPES (B). Active site residues complexed with HEPES in 1DG9 (Template)

### 2.4. Quality of model

The different aspects of structure of Dr-PTPs were validated using different tools. The stereochemical outcomes were analyzed using software PROCHECK where the restraints obtained were compared to the stereochemical properties of Bh-PTPs. The degree of violation of secondary structure elements were evaluated using ramachandran plot where 94% regions were found in allowed region and no dihedral region in disallowed regions confirms the validity of the model.

The 10 best structures of the Dr-PTPs were compared with Bh-PTPs crystal structure, both in free and complexed with HEPES. The RMSD based on backbone (α-carbon) were found 0.26 Å and 0.49 Å in the presence and absence of complexation (Table 1), respectively. These observations further confirmed the validity of the model beside the higher sequence identity. However, the conformational variability can be seen in several regions (1-6, 105-108, 134-136 and 121-123) due to presence of different amino acids inducing different orientation (Fig. 7a).

**Table 1.**

| Template | Target | DRMS |
| --- | --- | --- |
| Bovin heart PTPs (1BVH) with out inhibitor | Drosophila PTPs | 0.4840 |
| Bovin heart PTPs (1DG9) with inhibitor (HEPES) | Drosophila PTPs | 0.2549 |
| Bovin heart PTPs (1PNT) with inhibitor (PO4) | Drosophila PTPs | 0.1536 |

The fold energetic Dr-PTPs was calculated by program Prosa using template Bh-PTPs crystal structure. The comparison of energy was explored as shown by energy graph (Fig. 6).

**Figure 6.**
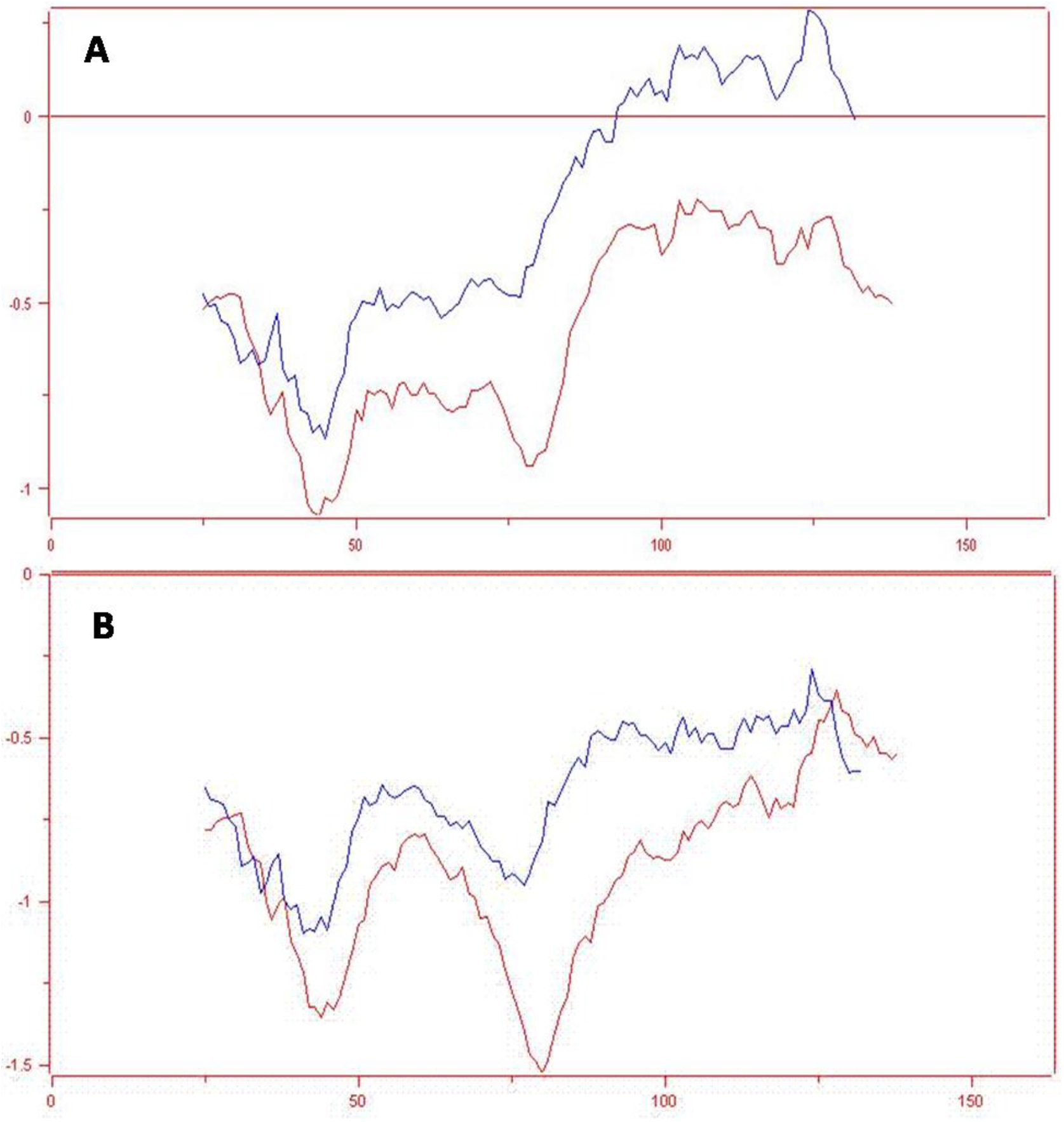
Comparison of PROSA II combined surface and pairing energy plots examined as a function of residue b/w template (blue) and the model (blue) for Bh-PTPs (A), 1DG9-PTPs, 1PNT-PTPs.

**Figure 7.**
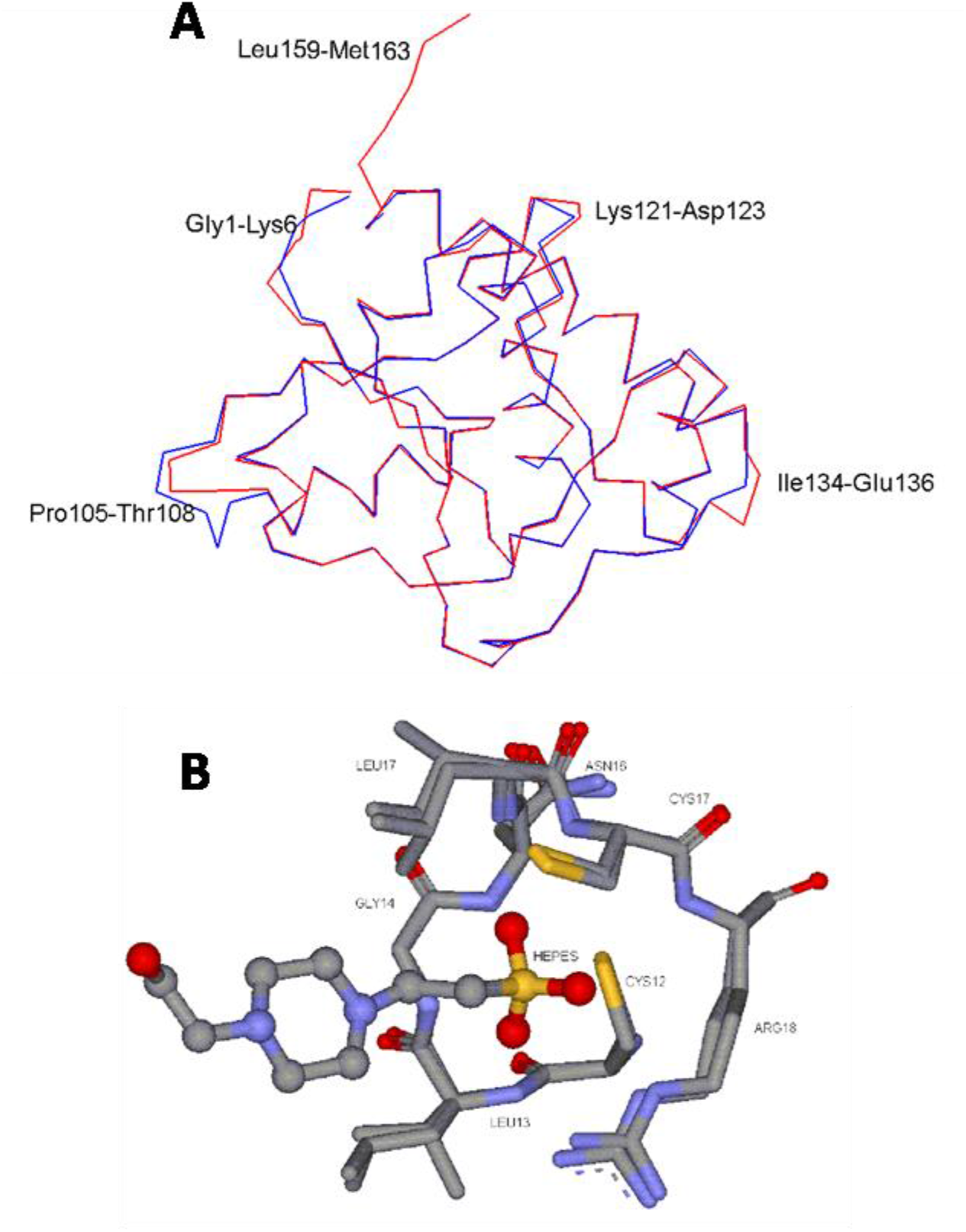
The active site is 100 % super imposed in template (Bh-PTPs) and Dr-PTPs. Active site is represented in sticks whereas inhibitor (HEPES) represents in ball and sticks style.

### 2.5. Active site and protein inhibitor interactions

The structural similarity of Dr-PTPs and Bh-PTPs and conformational resemblance of active site demonstrates the identity of catalytic mechanism. The catalytic reaction is triggered by the nucleophilic attack by Cys 3. The phosphate group at tyrosine group of the ligand or substrate bound at active site in orientation such that all the three oxygen atoms form hydrogen bonding with p-loop residues, making feasible the nucleophilic attack at phospho group by Cys3, Cys14, Cys16 and Cys18 where the Asp 129 works as proton donor resulting in the formation of phosphoenzyme intermediate (17, 19). The hydrolytic cleavage of intermediate takesplace resulting in the formation of inorganic phosphate.

The complexation of HEPES at active site of enzyme is stabilized by hydrophilic interactions like hydrogen bonding and hydrophobic electrostatic interactions and thus acts as a potent inhibitor. In Bh-PTPs (PDB: IDG9), the inhibitor HEPES is stabilized by forming seven hydrogen bonds with active site residues of enzyme. In similar fashion, nine hydrogen bonds were observed in Dr-PTPase. In case of Bh-PTPs, residues like Leu 13, Gly 14, Ile 16, Cys 17 and Arg 18 are involved in hydrogen bonding with inhibitor HEPES. Only three oxygens of HEPES involved in hydrogen bonding with active site residues of PTPs (Table 2). Residues like Val 14, Gly 15, Leu 17, Cys 18, Arg 19 participate in hydrogen bonding of Dr-PTPs (Target). Two nitrogen atoms and three oxygen atoms of HEPES (inhibitor) are involved in hydrogen bonding with active site residues of PTPase (Fig 7b).

**Table 2.** Ligand oxygen (HEPES)-Protein Nitrogen Bond length in PTPs complex

| Bovine Heart PTPs Complex with HEPES |  |  | Drosophila PTPs Complex with HEPES |  |  |
| --- | --- | --- | --- | --- | --- |
| 1DG9 | Amino acids | Distance (A<sup>0</sup>) | Model PTPs | Amino acids | Distance (A<sup>0</sup>) |
| EPE201:O1S | LEU13: N | 3.17 | EPE164:O1S | VA14: N | 3.18 |
| EPE201:O1S | GLY14: N | 2.98 | EPE164:O1S | GLY15: N | 3.02 |
| EPE201:O2S | ILE16: N | 3.04 | EPE164: C2 | LEU17: N | 3.06 |
| EPE201:O2S | CYS17: N | 2.88 | EPE164:O2S | CYS18: N | 2.88 |
| EPE201:O1S | ARG 18: NH2 | 2.93 | EPE164:O2S | ARG19: N | 3.26 |
| EPE201:O3S | ARG18: N | 3.2 | EPE164:O1S | ARG19: NH2 | 3.1 |
| EPE201:O3S | ARG18: NE | 2.94 | EPE164:O3S | ARG19: NE | 2.9 |
|  |  |  | EPE164: N1 | EPE164: N4 | 2.99 |
|  |  |  | EPE164: N4 | EPE164: N1 | 2.99 |

**Table 3.** Ligand oxygen (PO_4_)-Protein Nitrogen Bond length in PTPs complex

| Bovine Heart PTPs Complex with PO<sub>4</sub> |  |  | Drosophila PTPs Complex with PO<sub>4</sub> |  |  |
| --- | --- | --- | --- | --- | --- |
| 1PNT | Aminoacids | Distance((A) | PPD10 | Aminoacids | Distance(A) |
| PO4158:O3 | LEU13:N | 3.24 | PO4164:O3 | VAL14:N | 3.24 |
| PO4158:O3 | GLY14:N | 2.87 | PO4164:O3 | GLY15:N | 2.91 |
| PO4158:O2 | ILE16:N | 2.87 | PO4164:O2 | ASN16:N | 3.28 |
| PO4158:O2 | CYS17:N | 2.73 | PO4164:O2 | LEU17:N | 2.9 |
| PO4158:O3 | ARG18:NH2 | 2.92 | PO4164:O2 | CYS18:N | 2.74 |
| PO4158:O4 | ARG18:N | 3.29 | PO4164:O3 | ARG19:NH2 | 3.06 |
| PO4158:O4 | ARG18:NE | 2.78 | PO4164:O4 | ARG19:NE | 2.94 |
|  |  |  | PO4164:O1 | TYR131:OH | 3.17 |
|  |  |  | PO4164:O3 | PO4164:O4 | 2.51 |
|  |  |  | PO4164:O4 | PO4164:O3 | 2.51 |

The stabilization of HEPES for complexation with Bh-PTPs is favored by hydrophobic interaction by residues Ile 16 and Tyr 131, while in Dr-PTPs, Leu 17, His 51 and Tyr 131 are involved in hydrophobic interactions.

## 3. Materials and Methods

### 3.1. Sequence analysis

Sequence analysis of Dr–PTPs was obtained from SWISSPROT data base ^26^. The sequence homology from the protein data bank was obtained from BLAST ^27,28^, where modeller was used for target template alignment ^29^ for Dr–PTPs. The program Cluster was used for analysis of multiple sequences and adjustment of parameters made where necessary and finally, phylogenetic lineage was established with program phylip ^30,31^.

### 3.2. Model building and refinement

The three-dimensional model of Dr-PTPs was constructed using modeller (9V2) using Bh-PTPs crystal structure (PDB: 1DG9) as template model. The program was allowed to satisfy all dihedral angle, bond and spatial restraints and distances automatically as per default parameters. The input files consist of Dr-PTPs and Bh-PTPs aligned sequence. Several runs of calculations were performed to get more reliable and plausible model. The homology modeling was performed using standard parameters of calculations and known 3D structure models from protein data bank. The secondary structure elements of model were visualized using pymol and molmol ^16,31^ and other structural statistics was performed using psvs site (https://montelionelab.chem.rpi.edu/PSVS). The interaction of several ligands was analyzed using program Ligand Explorer (http://users.sdsc.edu/~q2hang/ligand)^32^.

### 3.3. Inhibitor modeling

The identification of hotspot residues, important for the target recognition and interaction was performed. The Dr-PTPs complexed with [N-(2-hydroxy ethyl) piperazine-N-2-ethanesulfonic acid sodium salt] (HEPES) was constructed using program modeller where the crystal structure 1DG9 was used as template. All these ligands known as potential inhibitors were models for the active sites for Dr-PTPs. The geometrical analysis, stereochemical analysis and all energies of bonds and dihedral restraints were analyzed. The homology models was subjected to program PROCHECK and ProSA for reliability of the model, secondary structure elements, backbone and energetic architecture and fold ^33–36^.

## 4. Conclusion

The sequence, secondary and tertiary structural similarities were studied in proteins Dr-PTPs and Bh-PTPs. The sequences of both proteins were found homologous for overall motifs and more especially for the active site motif represented by conserved residues C-(X)_5_-R. The comparative analysis of sequences is evident on the fact that this strong signature (CXXXXXR) at active site is the characteristic of low Molecular weight phosphotyrosine protein phosphatases. It was found that residues in 10-27, 81-88, and 127-130 regions were highly conserved with low Molecular weight PTPs. The structure obtained for this novel sequence were found of reliable and valid based on analysis performed by various protein structure validation tools as shown by the ramachandran plot, PROCHECK, energy of the fold and comparative analysis with other template crystal structures. The complexation profile of Dr-PTPs was also established based on the potent inhibitor HEPES. The overall stabilization factors important for the inhibitor complexation were studied and compared with known literature. We found that strong conformational similarity of Dr-PTPs with other homologous PTPs may shares same types of inhibitors exclusively considered to inhibit low Molecular weight phosphotyrosine protein PTPS. All these structural details obtained from the model are important for scheming further specific inhibitors. It can be used as an additional probe to decipher the discrete biological role of the low Molecular weight phosphotyrosine PTPS family and to explore the potential use of these macromolecular species as therapeutic targets.

## Data and Software Availability

The data acquired in this manuscript is available for method validation and reproducibility of the results. We used softwares Clustalx ^30,37^, Modeller^29^, Procheck^33^ for make sequence alignments and modeling studies. The data obtained is available in supporting information or otherwise mentioned in the articles.

## Acknowledgment

The project was financed by Lucian Blaga University of Sibiu and Hasso Plattner Foundation research grants LBUS-IRG-2021-07.

